# High-resolution genomic surveillance of 2014 ebolavirus using shared subclonal variants

**DOI:** 10.1101/013318

**Authors:** Kevin J. Emmett, Albert Lee, Hossein Khiabanian, Raul Rabadan

## Abstract

Viral outbreaks, such as the 2014 ebolavirus, can spread rapidly and have complex evolutionary dynamics, including coinfection and bulk transmission of multiple viral populations. Genomic surveillance can be hindered when the spread of the outbreak exceeds the evolutionary rate, in which case consensus approaches will have limited resolution. Deep sequencing of infected patients can identify genomic variants present in intrahost populations at subclonal frequencies (i.e. < 50%). Shared subclonal variants (SSVs) can provide additional phylogenetic resolution and inform about disease transmission patterns. Here, we use metrics from population genetics to analyze data from the 2014 ebolavirus outbreak in Sierra Leone and identify phylogenetic signal arising from SSVs. We use methods derived from information theory to measure a lower bound on transmission bottleneck size that is larger than one founder population, yet significantly smaller than the intrahost effective population. Our results demonstrate the important role of shared subclonal variants in genomic surveillance.

---

The West African ebolavirus outbreak arose in Guinea in late winter of 2014. As of December 10, 2014 there were 17,908 reported cases, with fatality rates as high as 75% in the most widely affected countries [1]. Despite the severity of the outbreak, genomic sampling of viral isolates has been limited [2–4]. Gire et al. [3] reported sequencing of ebolavirus samples from 78 patients collected during the initial introduction of the virus into Sierra Leone. Their analysis identified three distinct phylogenetic lineages and reported an evolutionary rate of ∼2 × 10^−3^ per site per year, or roughly one nucleotide change every two weeks. This corresponds to a rate almost twice as high as estimates from previous outbreaks [5]. These viral evolutionary dynamics in continuous human-to-human transmission are in agreement with measurements from intra-host evolution of filoviruses in primates [6].

Tracking the virus as it evolves during the course of the outbreak is an important goal of genomic surveillance, which has been facilitated with high-throughput genomics techniques. These techniques can rapidly sequence intrahost viral particles to high depth and provide estimates of allele frequencies at each position. Consensus-based approaches, which represent each patient with the majority allele measured at each position, have an inherent resolution of a single nucleotide. When the timescale of the outbreak is shorter than the average evolutionary time, as is the case with the 2014 ebolavirus outbreak, there will be insufficient genetic diversity for consensus approaches to provide good resolution into the evolution of the infectious agent. This suggests additional methods to gain what we term *subnucleotide resolution* into the evolutionary history of the outbreak.

Many samples in the Sierra Leone cohort were sequenced at sufficient depth to call subclonal variants, and initial analysis alluded to the presence of intrahost diversity and shared subclonal variants (SSVs) [3]. Based on genomic analysis as well as epidemiological data, Schieffelin et al. [4] reconstructed possible transmission chains; however, both studies missed evidence of SSVs that were later observed fixed in the cohort. Further, neither study attempted to integrate SSVs into a phylogenetic analysis, and while each suggested the presence of multiclonal transmissions, they did not attempt to assess the effective bottleneck size during possible direct transmissions or within the cohort.

Here, we incorporate subclonal diversity into a phylogenetic analysis using metrics from population genetics, following suggestions in Spielman et al. [7]. We identify several variants whose shared presence at subclonal frequencies shed light on interesting phylogenetic relationships not captured by consensus-based analyses. Finally, we introduce an information-theoretic method with which we estimate effective viral bottleneck size within a patient and during a transmission. When consensus diversity is limited and the outbreak spread exceeds the evolutionary rate, we show that measurement of subclonal diversity can provide valuable information for genomic surveillance.

## Subclonal variants

We performed variant calling on the Sierra Leone cohort from Gire et al. [3] using the 1976 Zaire isolate as reference. Because we were interested in confidently identifying SSVs, we retained only samples with mean sequencing depth ≥500×, which allowed us to obtain frequency estimates as low as 0.5% in 75 of the original 98 samples, from 64 of the original 78 patients. We identified an average of 568 variants per sample (range 559 to 582). Compared to the 1976 Zaire isolate, 541 variants were fixed in all Sierra Leone samples and 37 were found fixed during the current outbreak in at least one sample. We found 10 variants with frequencies ranging from ≥50% to <100%, and detected 221 subclonal variants at <50%, 45 of which were present in more than one sample. In our analysis, we identified all but two variants that Gire et al. reported, in addition to eight SSVs that they missed. Four of these variants were also found fixed in the cohort. Our analysis retained five patients with two, one patient with three, and one patient with four temporal samples.

## Phylogenetic patterns arising from SSVs

To incorporate the subnucleotide information from SSVs, we used Nei’s standard genetic distance [8], which assumes genetic differences are due to accumulation of mutations and genetic drift (Figure 1A). In patients with longitudinal data, we found limited variation in estimates of diversity during the course of the disease (Figure 1B), indicating the absence of hard selection sweeps and/or small population size effects after diagnosis. We also found stronger consensus-based similarity between pairs of samples with SSV compared to those without SSV (rank-sum test p-value: 5.1e-8), consistent with the expectation that samples sharing subclonal variants should be more related than those with no common subclonal variants (Figure 1C).

**Figure 1:**
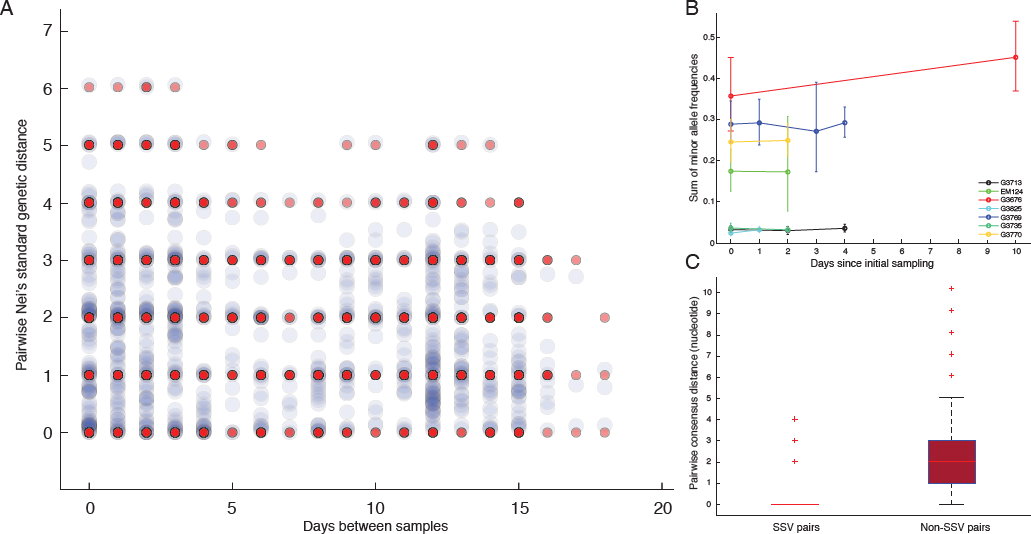
Subclonal variants. A) Consensus-based distances have inherent resolution of a single nucleotide; however, intrahost subclonal variants provide *subnucleotide resolution*. Red dots represent pairwise consensus distances and blue dots represent pairwise Nei’s genetic distances incorporating subclonal variants. (Here, we only show data collected from the chiefdom of Jawie.) B) There was minimal rise in intrahost genomic diversity during the course of the disease. The dip at the third temporal sample in G4769 corresponded to its lower sequencing depth and less sensitivity in identifying variants compared to patient’s other samples. The relative rise in diversity in G3676 corresponded to 1-2% change in frequency of six variants, still within their allele frequency confidence intervals. C) Samples that shared subclonal variants also had similar consensus genomes. In 26 pairs with SSV (Table 2), the mean pairwise consensus distance was significantly smaller than that of pairs with no SSV (<1 nucleotide in SSV pairs versus >2 in non-SSV pairs, rank-sum test p-value: 5.1e-8).

We used neighbor-joining to construct two phylogenetic trees: a consensus tree based on variants with frequencies ≥50%, and an SSV tree (Figure 2). Examining the consensus tree, we identified three clades, in agreement with previous analyses [3,9]. Four fixed variants (at sites 800, 8,928, 15,963, and 17,142) defined the split between clade 1 and clades 2 and 3. The split between clade 2 and clade 3 was due solely to a non-coding variant at position 10,218. Within each clade there were several degenerate sequences, and the finest resolution obtained was a single nucleotide. In this case, the sampling period spanned roughly one month, not sufficient time for substantial diversity to accumulate. When we considered the SSV tree, however, we observed that while the broad structure remained identical, the addition of subclonal information broke several degeneracies and introduced new branching patterns not observed in the consensus tree. We first noted the overwhelming role played by position 10,218 in generating the tree. We then highlighted notable substructures in the SSV tree and assigned them cluster labels. In particular, we identified seven clusters with phylogenetic relationships that arose due to the effect of SSVs (summarized in Table 1). Nonetheless, we do not suggest these patterns form persistent subclades, simply that they involve interesting patterns not discernible in a consensus analysis.

**Table 1:**
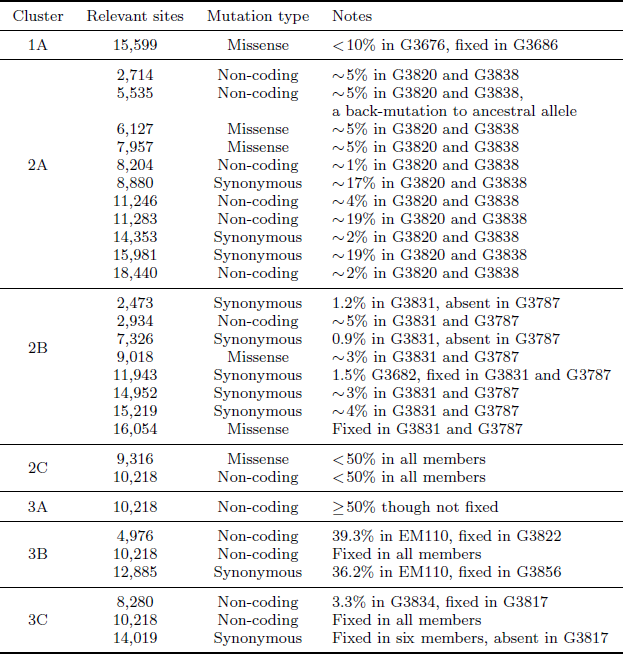
Summary of seven clusters with phylogenetic relationships that arose due to the effect of SSVs.

**Figure 2:**
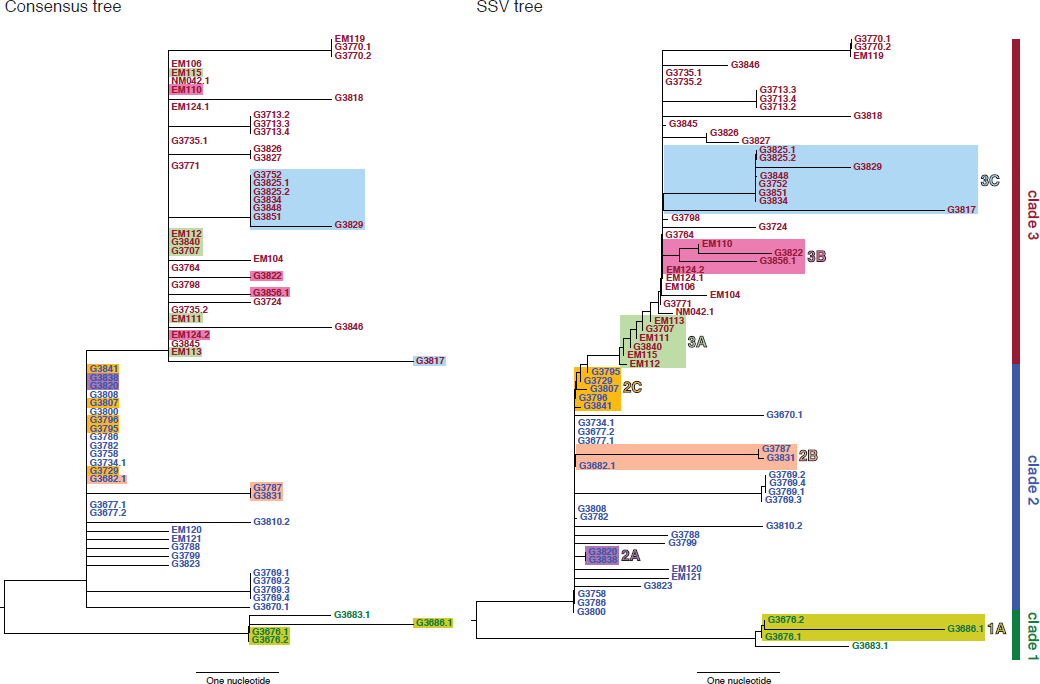
Consensus and SSV trees. The left panel shows a neighbor-joining tree constructed using the consensus sequence for each sample. The right panel shows a neighbor-joining tree constructed from incorporating SSVs and distances computed using Nei’s standard method. Specific differences between the two are highlighted and assigned cluster names within their respective clade.

In clade 1, a missense variant at site 15,599 rose from 7.5% in G3676 to become fixed in G3686. This defined cluster 1A. In clade 2, a synonymous variant at site 11,943 present at 1.5% in G3682 later became fixed in two patients, G3787 and G3831, constituting cluster 2A. A set of eleven shared variants between patients G3820 and G3838, including a possible back-mutation to the ancestral allele at position 5,535, defined cluster 2B. A set of samples that contained the variant at position 10,218 at <50%, as well as a unique SSV at position 9,316 (range 1.3% to 17.9%), defined cluster 2C.

In clade 3, cluster 3A was defined by the set of samples for which the variant at position 10,218 was present at >50% though not fixed. Cluster 3B represented an interesting relationship that was not visible in the consensus data. G3822 harbored a synonymous variant fixed at position 4,976 and G3856 harbored a synonymous variant fixed at position 12,885. In the consensus, no evidence existed to bring these two together, however we identified EM110, which harbored the variant alleles at both sites at <50%. This directly implicated EM110 as a putative ancestor of G3822 and G3856, with the two mutations existing unphased in EM110.

A non-coding variant at site 8,280, present at 3.3% in G3834, was found fixed in G3817. Because G3834 shared another fixed variant with other five patients, all seven patients were grouped together, defining cluster 3C. However, no phylogenetic relationship was consistent with this observed pattern, leading us to hypothesize either homoplasy or possible coinfection.

Finally, there were several SSVs that were shared between patients, but neighbor-joining did not bring together. A synonymous variant at site 10,509, present at only 1.3% in EM111, was observed at 100% in G3724. These two patients also harbored the variant at site 10,218 at 78% in EM111 and 100% in G3724, suggesting EM111 a plausible transmission ancestor of G3724. A shared subclonal variant at position 7,326 in G3831 (0.9%) and G3827 (6.25%) was the sole subclonal variant shared across clades 2 and 3. Again, homoplasy or coinfection are possible hypotheses to account for this pattern.

## Transmission bottleneck size and effective viral population

The presence of many shared subclonal variants between patients and the high fraction of reported cases in Sierra Leone from May 25th to June 20th represented by this dataset (∼70%) [10] strongly suggested direct transmission within this cohort [4]. However, comprehensive reconstruction of the exact chain of transmission has not been possible [4, 9]. Further, shared subclonal variants suggested *bulk* transmission, implying multiple viral populations can passage between individuals – more than a single founder population. While the idea of bulk transmission has been suggested in HIV [11,12], to our knowledge, estimates of transmission bottlenecks from deep sequencing have not been attempted. We therefore estimated the effective transmission bottleneck size within a patient and during transmission based on an information-theoretic method (Figure 3A). Here, we assumed minimal rise in diversity during the course of a patient’s disease, in agreement with our observation in seven patients for whom temporal samples were available (Figure 1B). In these patients, we estimated intrahost effective viral population size to be at least 10^5^ viral particles (range 260 to 1,183,885). In direct transmission amongst 16 pairs of patients, we found a lower bound of 10^2^ viral particles (range 0 to 800) effectively representing transmission population size, significantly smaller than the intrahost effective population (Figure 3B and Table 2).

**Table 2:** Summary of lower bounds on intrahost effective viral population and transmission bottleneck size.

| Sample 1 | Sample 2 | Type | Num. shared sites | $N_{\text{eff}}$ | $\sigma_{N_{\text{eff}}}$ |
| --- | --- | --- | --- | --- | --- |
| EM124.1 | EM124.2 | Intrahost | 7 | 405 | 153 |
| EM124.2 | EM124.1 | Intrahost | 5 | 260 | 116 |
| G3676.1 | G3676.2 | Intrahost | 3 | 327 | 189 |
| G3676.2 | G3676.1 | Intrahost | 6 | 626 | 256 |
| G3713.2 | G3713.3 | Intrahost | 2 | 8671 | 6131 |
| G3713.3 | G3713.2 | Intrahost | 2 | 8926 | 6312 |
| G3713.3 | G3713.4 | Intrahost | 2 | 2123 | 1501 |
| G3713.4 | G3713.3 | Intrahost | 2 | 2008 | 1420 |
| G3735.1 | G3735.2 | Intrahost | 4 | 2985 | 1492 |
| G3735.2 | G3735.1 | Intrahost | 3 | 2169 | 1252 |
| G3769.1 | G3769.2 | Intrahost | 3 | 1832 | 1058 |
| G3769.2 | G3769.1 | Intrahost | 3 | 1941 | 1121 |
| G3769.2 | G3769.3 | Intrahost | 3 | 598 | 345 |
| G3769.3 | G3769.2 | Intrahost | 3 | 625 | 361 |
| G3769.3 | G3769.4 | Intrahost | 3 | 672 | 388 |
| G3769.4 | G3769.3 | Intrahost | 3 | 643 | 371 |
| G3770.1 | G3770.2 | Intrahost | 9 | 5994 | 1998 |
| G3770.2 | G3770.1 | Intrahost | 10 | 6911 | 2185 |
| G3825.1 | G3825.2 | Intrahost | 1 | 593528 | 593528 |
| G3825.2 | G3825.1 | Intrahost | 2 | 1183885 | 837133 |
| G3827 | G3831 | Transmission | 15 | 107 | 28 |
| G3831 | G3827 | Transmission | 8 | 105 | 37 |
| G3826 | G3827 | Transmission | 5 | 4 | 2 |
| G3827 | G3826 | Transmission | 14 | 13 | 4 |
| G3820 | G3838 | Transmission | 12 | 750 | 217 |
| G3838 | G3820 | Transmission | 12 | 800 | 231 |
| G3817 | G3834 | Transmission | 5 | 1 | < 1 |
| G3834 | G3817 | Transmission | 3 | < 1 | < 1 |
| G3787 | G3831 | Transmission | 5 | 155 | 69 |
| G3831 | G3787 | Transmission | 6 | 154 | 63 |
| G3729 | G3795 | Transmission | 4 | 31 | 16 |
| G3795 | G3729 | Transmission | 14 | 92 | 25 |
| G3729 | G3796 | Transmission | 4 | 16 | 8 |
| G3796 | G3729 | Transmission | 7 | 46 | 17 |
| G3729 | G3807 | Transmission | 4 | 102 | 51 |
| G3807 | G3729 | Transmission | 9 | 263 | 88 |
| G3729 | G3841 | Transmission | 4 | 25 | 13 |
| G3841 | G3729 | Transmission | 8 | 68 | 24 |
| G3795 | G3796 | Transmission | 14 | 15 | 4 |
| G3796 | G3795 | Transmission | 7 | 15 | 6 |
| G3795 | G3807 | Transmission | 14 | 39 | 10 |
| G3807 | G3795 | Transmission | 9 | 34 | 11 |
| EM111 | EM115 | Transmission | 18 | 215 | 51 |
| EM115 | EM111 | Transmission | 8 | 89 | 31 |
| EM111 | G3724 | Transmission | 18 | 1 | < 1 |
| G3724 | EM111 | Transmission | 3 | < 1 | < 1 |
| G3676.1 | G3686.1 | Transmission | 3 | < 1 | < 1 |
| G3686.1 | G3676.1 | Transmission | 4 | 1 | < 1 |
| G3676.2 | G3686.1 | Transmission | 6 | 1 | < 1 |
| G3686.1 | G3676.2 | Transmission | 4 | 1 | < 1 |
| G3682.1 | G3787 | Transmission | 3 | < 1 | < 1 |
| G3787 | G3682.1 | Transmission | 7 | 1 | < 1 |

**Figure 3:**
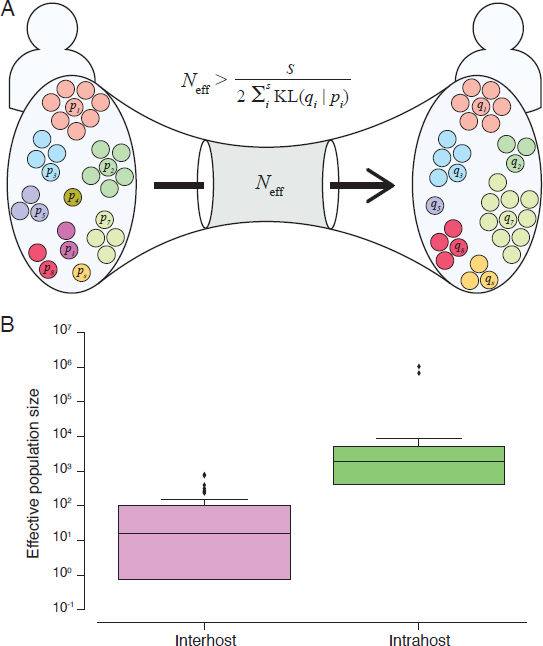
Effective transmission bottleneck size. A) Assuming minimal variation in variant frequency during the course of infection, we used binomial sampling to estimate lower bounds on effective transmission bottlenecks. SSVs provide evidence for bulk transmission of viral particles. This approach does not assume direct transmission, and is applicable whenever there is overlap of subclonal variants. B) Our results revealed a transmission bottleneck size significantly smaller than the intrahost effective viral population.

## Conclusion

Genomic surveillance promises to shed light on outbreak dynamics; however, when the rate of outbreak expansion exceeds the evolutionary rate, there may be insufficient resolution in consensus analysis to adequately track phylogenetic relationships. Advances in sequencing technologies now allow discovering very rare variants present as few as 1 out of 1,000 viral particles in each patient [6]. The study of viral evolutionary dynamics will benefit from treating patients as populations of viruses rather than a collection of single genomes. Our application of Nei’s standard genetic distance in reconstruction phylogenetic relationships incorporates population genetics methodologies. Tracking shared subclonal variants provides further information when tracking disease transmission and enhances resolution of evolutionary relationships to scales on the order of transmission time. Moreover, the information from SSV helps elucidate transmission processes beyond traditional consensus-based approaches, informing on the number of viral particles involved and potential sources of coinfections.

A caveat of our method is that it does not incorporate temporal information. Time in the case of 2014 ebolavirus outbreak can be a confounder, as the reported sampling time will not reflect where in the course of the infection a patient currently resides. This is of particularly important concern when the time scale of data collection is very short, on the same order as the infection period. Future work should focus on incorporating spatial and temporal annotations into the model.

## Methods

### Identifying shared subclonal variants

Raw sequencing reads from Gire et al. [3] were obtained from BioProject PRJNA257197. We mapped the reads to 1976 Zaire ebolavirus isolate (GenBank: NC_002549) using the Bowtie 2 aligner [13]. Because we were interested in calling shared subclonal variants, we limited our analysis to samples with mean coverage depth ≥500×. To identify statistically significant variants, we used the SAVI (Statistical Algorithm for Variant Identification) algorithm [14], which constructed empirical priors for the distribution of variant frequencies. From that prior, we obtained a corresponding high-credibility interval (posterior probability ≥1 − 10^−5^) for the frequency of each variant. Variants were considered present when observed with a lower bound frequency ≥0.5%. In some samples, due to higher sequencing depth, we were able to obtain frequency estimates below 0.5%; however, we chose this threshold to maintain consistent power for discovering variants at the cohort’s mean sequencing depth of 2,000× [15]. We filtered the variants for indel systematic sequencing errors mapping within homopolymeric tracts, and excluded adjacent, phased variants with highly correlated frequencies across all samples. We assessed variants’ strand bias against the dominant allele at their position using Fisher’s exact test. For patients with temporal samples, we excluded variants when the strand bias was significant in all samples (p-value ≤0.01). We applied a similar criterion to pairs of samples when calculating genetic distances and bottleneck sizes.

### Reconstructing phylogenetic trees

To compare viral populations between samples, we used Nei’s standard genetic distance [8]. If there existed *s* number of shared variants between samples 1 and 2,

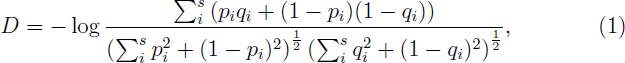

with *p*_*i*_ and *q*_*i*_ as the frequencies of variant *i* in samples 1 and 2, respectively. We constructed phylogenetic trees using the neighbor-joining method as implemented in PHYLIP [16]. Other genetic distances such as Nei’s *D*_*A*_ [17], Nei’s minimum genetic distance [18] produced similar results.

### Estimating effective viral population size

We assumed independence between variants and minimal variation in their frequencies during the course of a patient’s disease, consistent with the collected data, and estimated a lower bound on transmission bottleneck size with methods derived from information theory (Figure 3A). We applied a similar approach to estimate effective population size within a patient when temporal data were available.

If there existed *n*_*i*_ copies of virus harboring variant *i* in sample 1, the probability of observing *m*_*i*_ viral particles with the same variant in sample 2 could be described with binomial sampling as

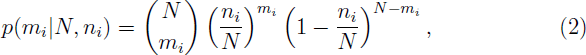

with *N* as the bottleneck size. After changing the variable *n*_*i*_ to *Np*_*i*_ and *m*_*i*_ to *Nq*_*i*_, respectively, for *s* number of shared variants, the likelihood of the observed state would become

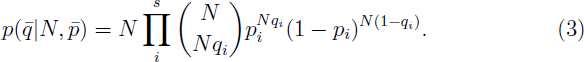

We then followed Stirling’s approximation for factorials, 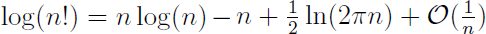, and derived the log-likelihood to be

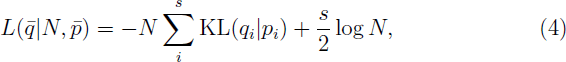

with KL(*q*_*i*_∣*p*_*i*_) representing the Kullback-Leibler divergence of *q*_*i*_ from *p*_*i*_ [19]. Therefore, the maximum likelihood estimate of *N*, describing the lower bound on effective bottleneck size, was

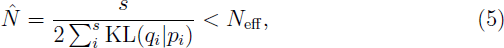

with variance

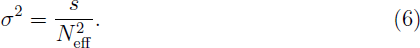

We calculated effective bottleneck size for pairs of patients who shared at least one non-fixed variant, excluding the variant at site 10,218. We calculated *N*_eff_ in both orientations, from sample 1 to sample 2 and vice versa. We would like to emphasize that this is a lower bound on *N*_eff_ as sequencing of viral populations in each sample should be considered as additional Markov processes.

## Acknowledgments

We thank Marta Luksza, Daniel Rosenbloom, and Ohad Balaga for helpful discussions. This work was supported by NIH Grant U54 CA121852 and the Defense Threat Reduction Agency (DTRA) Project HDTRA1-14-1-0016.

## References

1 CDC (2014) Update: Ebola Virus Disease Epidemic – West Africa, December 2014. Morbidity and Mortality Weekly Report 63: 1199–1201.

2 Baize S, Pannetier D, Oestereich L, Rieger T, Koivogui L, et al. (2014) Emergence of zaire ebola virus disease in guinea. New England Journal of Medicine 371: 1418–25.

3 Gire SK, Goba A, Andersen KG, Sealfon RS, Park DJ, et al. (2014) Genomic surveillance elucidates Ebola virus origin and transmission during the 2014 outbreak. Science 345: 1369–72.

4 Schieffelin JS, Shaffer JG, Goba A, Gbakie M, Gire SK, et al. (2014) Clinical illness and outcomes in patients with Ebola in Sierra Leone. New England Journal of Medicine 371: 2092–100.

5 Carroll SA, Towner JS, Sealy TK, McMullan LK, Khristova ML, et al. (2013) Molecular Evolution of Viruses of the Family Filoviridae Based on 97 Whole-Genome Sequences. Journal of Virology 87: 2608–2616.

6 Khiabanian H, Carpenter Z, Kugelman J, Chan J, Trifonov V, et al. (2014) Viral diversity and clonal evolution from unphased genomic data. BMC Genomics 15.

7 Spielman SJ, Meyer AG, Wilke CO (2014) Increased evolutionary rate in the 2014 West African Ebola outbreak is due to transient polymorphism and not positive selection. bioRxiv.

8 Nei M (1972) Genetic distance between populations. American Naturalist 106: 283.

9 Luksza M, Bedford T, Lassig M (2014) Epidemiological and evolutionary analysis of the 2014 ebola virus outbreak. arXiv:14111722.

10 Dixon MG, Schafer I (2014) Ebola viral disease outbreak – west africa, 2014. Morbidity and Mortality Weekly Report 63: 548–51.

11 Keele BF, Giorgi EE, Salazar-Gonzalez JF, Decker JM, Pham KT, et al. (2008) Identification and characterization of transmitted and early founder virus envelopes in primary HIV-1 infection. Proceedings of the National Academy of Sciences 105: 7552–7557.

12 Carlson J, Schaefer M, Monaco D, Batorsky R, Claiborne D, et al. (2014) Selection bias at the heterosexual HIV-1 transmission bottleneck. Science 345.

13 Langmead B, Salzberg SL (2012) Fast gapped-read alignment with Bowtie 2. Nature Methods 9: 357–9.

14 Trifonov V, Pasqualucci L, Tiacci E, Falini B, Rabadan R (2013) SAVI: a statistical algorithm for variant frequency identification. BMC Systems Biology 7 Suppl 2: S2.

15 Rossi D, Khiabanian H, Spina V, Ciardullo C, Bruscaggin A, et al. (2014) Clinical impact of small TP53 mutated subclones in chronic lymphocytic leukemia. Blood 123: 2139–47.

16 Felsenstein J, Nei M (1989) PHYLIP - Phylogeny Inference Package (Version 3.2). Cladistics 5: 164–166.

17 Nei M, Tajima F Y T (1983) Accuracy of estimated phylogenetic trees from molecular data. ii. gene frequency data. Journal of Molecular Evolution 19: 153–170.

18 Nei M, Roychoudhury AK (1974) Genic variation within and between the three major races of man, Caucasoids, Negroids, and Mongoloids. The American Journal of Human Genetics 26: 421–443.

19 Kullback S, Leibler RA (1951) On Information and Sufficiency. Annals of Mathematical Statistics 22: 142–143.

